# Duration reproduction under memory pressure: Modeling the roles of visual memory load in duration encoding and reproduction

**DOI:** 10.1101/2022.02.10.479853

**Authors:** Xuelian Zang, Xiuna Zhu, Fredrik Allenmark, Jiao Wu, Hermann J. Müller, Stefan Glasauer, Zhuanghua Shi

**Affiliations:** Center for Cognition and Brain Disorders, Affiliated Hospital of Hangzhou Normal University, 310015, China; General and Experimental Psychology, Department of Psychology, LMU Munich, 80802, Germany; Computational Neuroscience, Institute for Medical Technology, Brandenburg Technical University Cottbus-Senftenberg, 03046, Cottbus, Germany

**Keywords:** time perception, dual-task performance, attention-sharing, cognitive/memory load, Bayesian integration, central-tendency effect

## Abstract

Duration estimates are often biased by the sampled statistical context, yielding the classical central-tendency effect, i.e., short durations are over- and long duration underestimated. Most studies of the central-tendency bias have primarily focused on the integration of the sensory measure and the prior information, without considering any cognitive limits. Here, we investigated the impact of cognitive (visual working-memory) load on duration estimation in the duration encoding and reproduction stages. In four experiments, observers had to perform a dual, attention-sharing task: reproducing a given duration (primary) and memorizing a variable set of color patches (secondary). We found an increase in memory load (i.e., set size) during the duration-encoding stage to increase the central-tendency bias, while shortening the reproduced duration in general; in contrast, increasing the load during the reproduction stage prolonged the reproduced duration, without influencing the central tendency. By integrating an attentional-sharing account into a hierarchical Bayesian model, we were able to predict both the general over- and underestimation and the central-tendency effects observed in all four experiments. The model suggests that memory pressure during the encoding stage increases the sensory noise, which elevates the central-tendency effect. In contrast, memory pressure during the reproduction stage only influences the monitoring of elapsed time, leading to a general duration over-reproduction without impacting the central tendency.

## Introduction

Accurate timing is essential for proper actions in our daily activities, such as synchronizing our body movements to a rhythm in music or pronouncing subtly different syllables such as /pa/ and /ba/ with voice-onset time (VOT) in the millisecond range. Yet, subjective time never stops surprising us. Most of us have conscious experiences of situations where time flies or time drags. More surprisingly, distortion of time often happens without us explicitly knowing it. One classical example of such implicit time distortion is the Vierordt effect (better known as the central-tendency effect), reported a century and a half ago (Vierordt, 1868), which describes the phenomenon of short intervals being overestimated and long intervals underestimated (Glasauer & Shi, 2021a, 2021b; Jazayeri & Shadlen, 2010; Lejeune & Wearden, 2009; Shi et al., 2013). Notably, central-tendency effects are ubiquitous in different types of sensory magnitude estimation (Petzschner et al., 2015), such as in spatial distance and angular (rotational-body) displacement judgments (Petzschner et al., 2012; Petzschner & Glasauer, 2011; Teghtsoonian & Teghtsoonian, 1978).

A common explanation of the central-tendency effect is that magnitude estimation is not only based on the sensory measurement but also influenced by past experience, in particular, of the range and distribution of tested intervals. According to the view of Bayesian inference, the brain integrates the sensory measure and prior knowledge together to boost the precision of the estimation – which, while being beneficial in most cases, also engenders a byproduct: a central-tendency bias (for reviews, see Petzschner et al., 2015; Shi et al., 2013). Optimal integration of the sensory input and prior knowledge depends on their respective reliability, measured by the inverse of their variance (the precision). When the sensory measurement has high precision, such as in professional drummers, there would be less influence of prior knowledge, resulting in a lesser central-tendency bias (Cicchini et al., 2012). Importantly, the integration of the sensory measure and the prior is likely involved in the working memory (WM), so that the cognitive load (the demands on WM capacity) may impact both the sensory estimate and the prior representation in terms of their means and variances. However, the role of WM on Bayesian inference of time perception has been largely neglected in the literature.

Although not focusing on Bayesian inference of time perception, Fortin and Rousseau (1998) reported two separable effects of cognitive load on duration encoding and duration reproduction, respectively. In their dual-task design, the secondary task was a Sternberg memory task with a memory set of 1, 3, or 6 digits, presented successively prior to the primary temporal reproduction task, where the latter consisted (on a given trial) of an initial duration-production phase (with two beeps demarcating a duration) followed by the duration-reproduction phase (two taps generated by the participants). The memory probe (a digit that was or was not part of the memory set) was shown either during the duration-production or the -reproduction phase, and the response to the memory task (either positive or negative) was to be issued using the key of the first (for probes presented during the production phase) or, respectively, the second (for probes presented during the reproduction phase) tap of the temporal reproduction response (one of two keys). Fortin and Rousseau found the reproduced duration to be shortened when the memory probe was presented during the production phase, but lengthened when it was shown during the reproduction phase. They took this finding to support an attention-sharing account (Fortin & Rousseau, 1998; Macar et al., 1994), according to which attentional resources are shared between the timing process and other, non-temporal cognitive processes. When attention is diverted away from the primary task by other concurrent, non-temporal processes in the temporal encoding (i.e., production) phase, the perceived duration is shortened. On the other hand, if the non-temporal process interferes with the reproduction of a given duration, the lapse in the monitoring of the passage of time will lengthen the reproduced duration. Similar findings have been reported in other timing studies (e.g., Fortin & Couture, 2002; Fortin & Massé, 2000), as well as in a study of non-temporal magnitude estimation(Glasauer et al., 2007).

It should be noted that the above studies focused on the over- and under-estimation caused by the memory load. Thus, it remains unclear how memory load influences the uncertainty of the magnitude encoding and prior representation, as well as how it impacts the subsequent magnitude reproduction. Interestingly, a recent study (Allred et al., 2016) on working memory and spatial-length judgments suggests that high cognitive load induced by the secondary working memory task could lead to a coarser memory representation of spatial length, that is, high uncertainty, yielding a strong central-tendency effect. However, in (Allred et al., 2016) design, the secondary working-memory task extended across the whole spatial-length judgment task and so did not permit dissociating between load influences on the (length) encoding vs. the (length) reproduction stages. Accordingly, whether the central tendency would be differentially influenced by the cognitive load on the encoding and reproduction phases (in duration judgments) remains unclear.

Taking together the literature reviewed above, we hypothesized that cognitive load would influence both the perceived and reproduced durations, as well as the variability of the estimates, which would further affect Bayesian inference in time estimation (Jazayeri & Shadlen, 2010; Shi et al., 2013; Shi & Burr, 2016). Specifically, we expected increasing cognitive load during the sensory encoding stage not only to lead to a general underestimation of the duration (in line with Fortin & Rousseau, 1998), but also to decrease the reliability of the estimate. Accordingly, Bayesian integration of the sensory measure and the prior (also referred to as “memory mixing” in B. M. Gu & Meck, 2011; Jazayeri & Shadlen, 2010) would predict higher cognitive load to engender a stronger central-tendency bias. By contrast, introducing cognitive load during the reproduction stage would lengthen the reproduced duration (i.e., produce a general overestimation), and likely also increase the variability of the reproduction. However, given that no additional cognitive load is imposed on the sensory encoding stage, the reliability of the sensory estimate, and thus the Bayesian integration, would be unaffected by the cognitive load introduced during duration reproduction. When cognitive load remains high during both the duration production and reproduction phases, the underestimate from the production and the overestimate from the reproduction may cancel each other, at least to some extent and so we may not be able to observe a general bias. However, increasing the uncertainty in the sensory representation may cause a stronger central-tendency effect – a similar pattern to that recently reported in a non-temporal task(Allred et al., 2016).

To test these hypotheses, we adopted a dual-task paradigm, consisting of a secondary *visual* working-memory task (with low, medium, or high load) and a primary duration production-reproduction task, in four experiments (see Table 1). Importantly, we manipulated the ‘spanning’ of the secondary task – that is, the period over which the memory-set items had to be maintained – in relation to the primary timing task in such a way that the memory task influenced different stages (production, reproduction, or both) of the timing task. Specifically, in Experiment 1, the memory task was introduced after the timing task, providing a baseline. In Experiment 2, the memory task spanned the duration-production phase, to examine the impact of cognitive load on duration encoding. In Experiment 3, by contrast, it spanned the duration-reproduction phase to examine the impact of cognitive load on the duration reproduction. Finally, in Experiment 4, the memory task extended over the whole timing task (i.e., both the production and reproduction phases), to examine the combined effect of a non-temporal task on the timing task, while also serving as a comparison to the study of memory load on the central tendency in a spatial task (Allred et al., 2016).

**Table 1.** Experimental designs with the dual tasks

| Experiment | Interleaved order of the dual task |  |  |  |
| --- | --- | --- | --- | --- |
| 1 | Production | Reproduction | Memory phase | Memory test phase |
| 2 | Memory phase | Production | Memory test phase | Reproduction |
| 3 | Production | Memory phase | Reproduction | Memory test phase |
| 4 | Memory phase | Production | Reproduction | Memory test phase |

Given that the attentional sharing account (Fortin & Rousseau, 1998; Macar et al., 1994) makes clear predictions of how memory load would influence the production and reproduction stages, and Bayesian inference makes quantitative predictions of how the sensory estimate is integrated with the prior, we also developed a general computational framework for duration encoding, Bayesian integration and duration reproduction, taking into consideration possible influences of cognitive load, based on our behavioral findings. We identified the role of attentional sharing and the temporal stage of prior integration in time estimation. Specifically, owing to attentional sharing between the concurrent working-memory task and duration estimation, participants displayed underestimation with memory pressure during the production, but overestimation in the reproduction phase. These two opposing influences canceled each other, diminishing the general bias when memory pressure covers both the production and reproduction phases. Moreover, we found an increased central-tendency effect with memory pressure during the production phase, but no significant changes with memory pressure during the reproduction phase.

## Methods

### Participants

Different groups, each of 16 volunteers (23-34 years old), were recruited for each experiment (9 females in Experiments 1, 2, and 4, 8 females in Experiment 3); all of them had self-reported normal or corrected-to-normal vision and normal hearing. The sample size was determined based on (Fortin & Rousseau, 1998) study, in which ten participants yielded significant under- and over-estimations. On the conservative side, we recruited 16 participants. All participants were naive as to the purpose of the experiments and received 9 Euro per hour for their service. The experiment was approved by the Ethics Committee of the Department of Psychology of LMU Munich.

### Apparatus

The experiments were conducted in a sound-isolated, dimly lit cabin (5.24 cd/m^2^). The visual stimuli were presented on a 21” LACIE CRT monitor, with a refresh rate of 100 Hz. The viewing distance was fixed to 57 cm (maintained by the use of a chin rest). The experimental program was developed using Matlab (Mathworks Inc.) and Psychtoolbox (Kleiner et al., 2007).

### Stimuli and tasks

The experiments were dual-task experiments, consisting of a duration production-reproduction task and a visual working-memory task on each trial.

The ***duration production-reproduction task*** (see Figure 1 for an example) consisted of two phases: In the first, production phase, a grey disk (36.5 cd/m^2^, 4.15° in diameter) was presented in the center of the monitor (on a dark background: 16.7 cd/m^2^) for a given duration, randomly sampled from 500, 800, 1100, 1400, or 1700 ms. Participants were instructed to encode and retain the duration of the grey disk. In the following reproduction phase, participants were asked to reproduce the perceived duration of the grey disk as accurately as possible by pressing and holding the down-arrow key. The key-press triggered a visual display with a grey disk, which stayed on the screen until the key was released. Participants were asked to reproduce, as accurately as possible, the (presentation) duration of the grey disk from the production phase.

**Figure 1.**
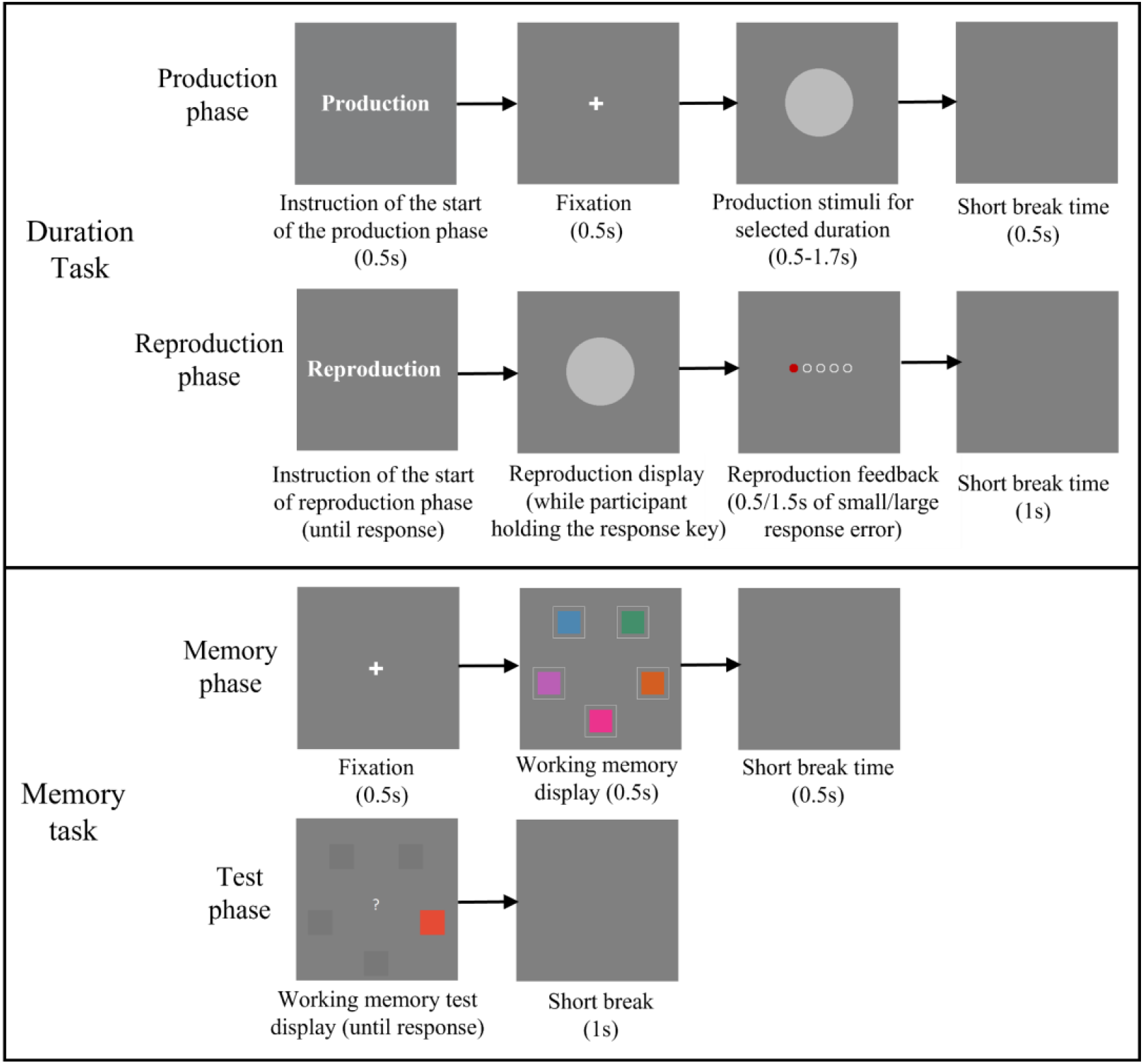
Schematic illustration of the dual tasks used in Experiment 1. The duration task includes the production and reproduction phases (depicted in the upper two panels). The working memory task also includes two phases: the memory phase and the test phase (depicted in the lower two panels). In the example, the correct answer would have been “different”.

The ***visual working-memory task*** also consisted of two phases: a memory phase and a test phase (see Figure 1). During the memory phase, a number (randomly selected 1, 3, or 5) of squares were presented on an invisible circle (radius approximately 4.15°) on the dark background. Each square (subtending 2.14° × 2.14°) was filled with a color randomly selected from 180 color values uniformly distributed along a circle in CIE 1976 (L* = 70) color space (van den Berg et al., 2012; Zhang & Luck, 2008). Adjacent items were arranged equidistantly in displays with three or, respectively, five items; in one-item displays, the single item appeared always at the bottom position on the invisible circle. Participants had to encode and retain the color in which a square (at a given location) appeared (with the memory load increasing with the number of squares). In the test phase, one item location from the memory phase was selected as the location of the test (or probe) stimulus. The color of the probe square was either the same or different (randomly and equally determined) relative to the previous (memory) item at that location. In case of a “different” probe, the color was randomly selected from the remaining 179 possible colors. [Note that the probe square was presented without the thin grey outline that framed the items in the memory phase (see Figure 1).] Participants were asked to indicate whether the probe item had the same or a different color as the previous item at that location by pressing either the left (same) or right (different) arrow key.

### Design and procedure

#### Experiment 1

served as a baseline. Participants were presented with the two tasks successively: the duration task first and the memory task second. Thus, any interference between the two tasks would have been minimal (see Table 1). The experiment consisted of 18 blocks of 20 trials each. The working-memory load (1, 3, or 5 items) was fixed per block, but randomly counter-balanced across the 18 blocks. Using a block-design for the memory load in this baseline experiment was meant to rule out any possible interference of the memory load on the updating of the duration prior across trials.

The duration task started with the cue word ‘Production’ on the screen for 500 ms, indicating the *production phase*. Then, a fixation dot appeared for 500 ms, followed by a grey disk in the center of the monitor; this remained visible for a given trial duration (0.5 to 1.7 s), after which the display turned back to a blank screen for 500 ms. Next, the second cue display with the word ‘Reproduction’ appeared, prompting the start of the *reproduction phase*. Participants followed their own pace to initiate the reproduction, which required them to press and hold down the down-arrow key. Immediately after the key press, a grey disk appeared and remained visible until the key was released (i.e., the visible disk duration served as the reproduced event). The duration of the key pressing was recorded as the reproduced duration. After the participant released the response key, a feedback display was presented showing the relative reproduction error (i.e., the ratio of the reproduction error to the given trial duration) by highlighting one of five linearly arranged dots (see Figure 1). The five dots, from the left to the right, were mapped to the relative error ranges: below −30%, between [-30%,-5%], [-5%, 5%], [5%, 30%] and greater than 30%, respectively. The three central dots were colored green, and the left- and right-most dots red, indicating how large a reproduction error was made. When the relative error was between −30% and 30%, the error feedback display was presented for 500 ms; otherwise, it was presented for 1500 ms, alerting participants that the error was too high.

After a break of 1 s with a blank screen, the working-memory task started with a fixation cross presented for 500 ms, followed by a memory display containing one, three, or five colored squares visible for 500 ms (*memory phase*). After a blank screen of 500 ms, a probe display with a single color patch at one of the previous (i.e., memory-phase item) locations (*test phase*). Participants had to indicate whether this probe patch was colored the same vs. differently relative to the (coincident) item in the memory display, by pressing the left (‘same’) or the right (‘different’) arrow key. After a one second interval, the next trial began.

#### Experiment 2

examined whether maintaining information in working memory during the production (but not the reproduction) phase would affect the sensory duration measurement and further influence the use or updating of priors. The tasks and displays were essentially the same as in Experiment 1, except that, in the trial-event sequence, the working-memory task spanned the duration production (and not the reproduction) phase. That is, each trial started with the *memory phase*, followed by the duration *production phase*. Next, participants performed the *memory test*, before proceeding to the duration *reproduction phase* (see Table 1 and Figure 1). Given that the prior updating was not impacted by memory load in the baseline experiment (see Results of Experiment 1), we adopted a trial-wise design for the memory load in Experiments 2 to 4, effectively making the load unpredictable; accordingly, the influence of memory load (if any) would be locally trial-based for each tested duration.

#### Experiment 3

was essentially the same as Experiment 2, except that now the working-memory task spanned the reproduction (rather than the production) phase. That is, each trial started first with the *production phase*, followed by the *memory phase*; then, participants had to *reproduce* the given (trial) duration and finally perform the *memory test* (see Table 1 and Figure 1).

#### Experiment 4

examined how duration estimation would be affected by maintaining the working-memory information across both the production and reproduction phases. That is, the task sequence on each trial was: memory phase, duration-production phase, duration-reproduction phase, and finally memory-test phase (see Table 1).

## Modeling of duration estimation under memory load

The duration production-reproduction task involves the component stages of duration encoding, Bayesian integration, and duration reproduction. Here we proposed a generative processing architecture and potential influences of the memory load on these stages.

### 1. Duration encoding

We assume that, while the ‘raw’ sensory measure (*S*) of given sample duration (D) is not influenced by cognitive load, its representation in working memory *S_wm_* may be affected by the load. Given that the scalar property (i.e., Weber scaling) is the key feature of duration estimation (Gibbon et al., 1984; Shi et al., 2013), we further assume logarithmic scaling of the sensory measure (*S*), to simplify calculation; that is, 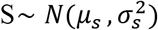, where *μ_s_* is the mean of the logarithmic scale representation of the given sample interval D (see Petzschner & Glasauer, 2011; Roach et al., 2017), and 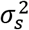 reflects the variance of internal-measurement noise (*ϵ*):

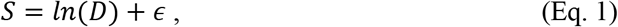

According to the classical internal-clock and attentional-gate models (Block & Zakay, 1997; Gibbon et al., 1984), sharing of attention by a concurrent non-temporal task would lead to an extra loss of the accumulated clock ticks, as well as increasing the noise of the memory representation of the sensory measurement (***S_wm_***). Here, for modeling the concurrent memory-load effects, we assume – as a simple approximation and following the principle of ‘Occam’s razor’ – that both the number of ticks and the noise modulation (as represented on the logarithmic scale) are *linearly* affected by the memory load.^1^ That is, the memory representation is normally distributed, 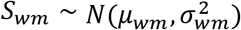, with both the mean *μ_wm_* and the variance 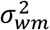 influenced by the memory load linearly:

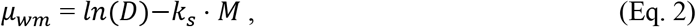

and

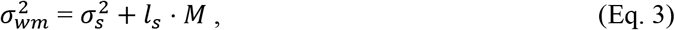

where *M* represents the level of the working-memory load (set to 1, 2, and 3 for the low, medium, and high load, respectively^2^), and *k_s_* and *l_s_* are scaling factors of the mean and the variance of the memory representation, respectively. Note that based on the attentional-gate model, we specifically assume that the cognitive-load influence on duration encoding in Eqs. 2 and 3, *k_s_* and *l_s_*, are constrained to be non-negative (*k_s_* ≥ 0, *l_s_* ≥ 0). Given that the duration-production phase was *non*-overlapping with the secondary working-memory task in Experiments 1 and 3, *k_s_* and *l_s_* were set to zero for those experiments (i.e., there would be no influence of the secondary task on the production phase). In more detail, *k_s_* · *M* represents the loss of clock ticks during the accumulation process. In addition, the cognitive load increases the variance of the memory representation in a linear fashion (Bays, 2015), which is captured by the term *l_s_* · *M*.

### 2. Bayesian integration

The classical central-tendency effect shown in duration reproduction can be explained by memory mixing between the internal prior of the sampled durations and the sensory measure (Acerbi et al., 2012; B.-M. Gu et al., 2016; Jazayeri & Shadlen, 2010; Penney et al., 2000). Given that the sampled durations were the same across the three memory-load conditions, we assume the internal prior was the same for the different memory loads, following the normal distribution with the mean *μ_p_* and the variance 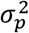 (both parameters in logarithmic space). The internal prior is then integrated with the memory representation of duration *S_wm_* according to the Bayes rule, which minimizes the uncertainty of the final duration representation. According to Bayesian inference, the posterior distribution 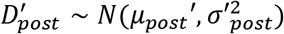 of the final memory representation can be estimated by

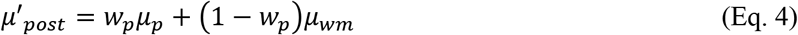

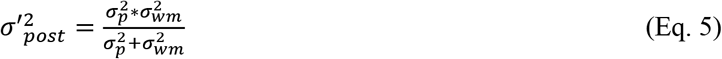

The optimal weight *w_p_* is proportional to the inverse variability of the priors 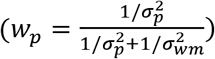, which indicates the relative influence of the prior. The weight plays a key role in the *l^/^ōp+^1/^σ_wm_* central-tendency effect (Jazayeri & Shadlen, 2010; Shi et al., 2013): larger values of *w_p_* mean a stronger central-tendency effect.

### 3. Duration reproduction

Given that attention is required for the time-monitoring during the reproduction stage, the memory load might influence the final reproduction output. Our model assumes that the memory load influences the monitoring of the elapsed time of the reproduction, that is: there is a continuous monitoring of the elapsed time starting from the key press (*μ_elapsed_*) and comparison of the elapsed time with the duration held in memory (*μ′_post_*). Similar to the production stage, the sensory representation of the elapsed time can be distorted by the memory load. Accordingly, the comparison is:

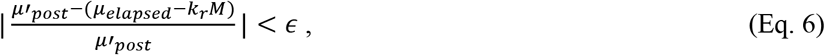

which is equivalent to comparing the elapsed time to the mean duration *μ′_post_* + *k_r_M*. In order to compare the model calculations to the observed reproduction behavior, we transferred the logarithmic space representation in the model back to the linear space, using the lognormal distribution with mean and variance:

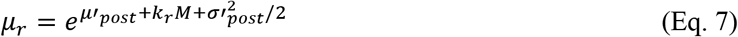

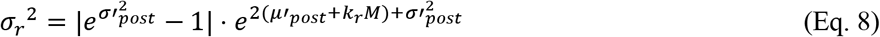

In addition, we assume the uncertainty of reproduction is further influenced by motor noise 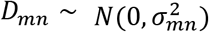, but with the the influence of the motor noise decreasing as the duration increases:

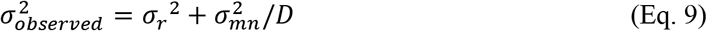

Thus, we have seven parameters in total in the above model framework: the memory load scaling parameters of the mean and variance of the duration encoding (*k_s_*, *l_s_*), the variance of sensory measurement 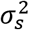, the memory-load scaling factor *k_r_* in the reproduction phase, the motor noise *σ*^2^_*mn*_, and the mean and variance of the prior (*μ_p_*, *σ*_*p*_^2^).

Moreover, the general over-/under-estimation is usually measured by the average mean reproduction, which has a positive linear relation to an indicator called the indifferent point (IP), i.e.: the point at which the duration reproduction is veridical (Lejeune & Wearden, 2009)^3^. Based on Eqs. 2, 4, and 7, and letting the reproduced duration equal to the given duration (by the definition of IP), the log of the indifference point can be written as

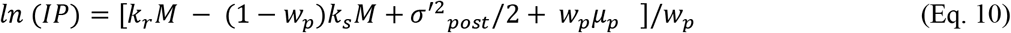

As we can see from Eq. 10, the first-order impact of the memory load is the combination of the scaling factors *k_s_* and *k_r_*, and the second-order impact is mediated through the variance (*μ_p_* and *w_p_*).

### 4. Parameter estimation

We adopted Stan, a platform for statistical modeling and Bayesian statistical inference (Bürkner, 2016; Stan Development Team, 2020), to estimate the parameters in the hierarchical Bayesian modeling. To complete the Bayesian hierarchical model, we have used standard non-informative priors on those hyperparameters (see Figure 2). In our proposed model, the seven parameters θ =(*k_s_*, *σ_s_*, *l_s_*, *μ_p_*, *σ_p_*, *k_r_*, *σ_mn_*) are sampled from their respective prior distributions p(θ) with hyperparameters. The distributional belief about parameters θ can be denoted as a conditional probability function *p*(*θ*|*D_observed_*). Using the Bayes rule, the posterior distribution *p*(*θ*|*D_observed_*) ∝ *p*(*θ*) × *p*(*D_observed_*|*θ*) can be derived from the prior distribution, p(θ), and a likelihood, *p*(*D_observed_*|*θ*).

**Figure 2.**
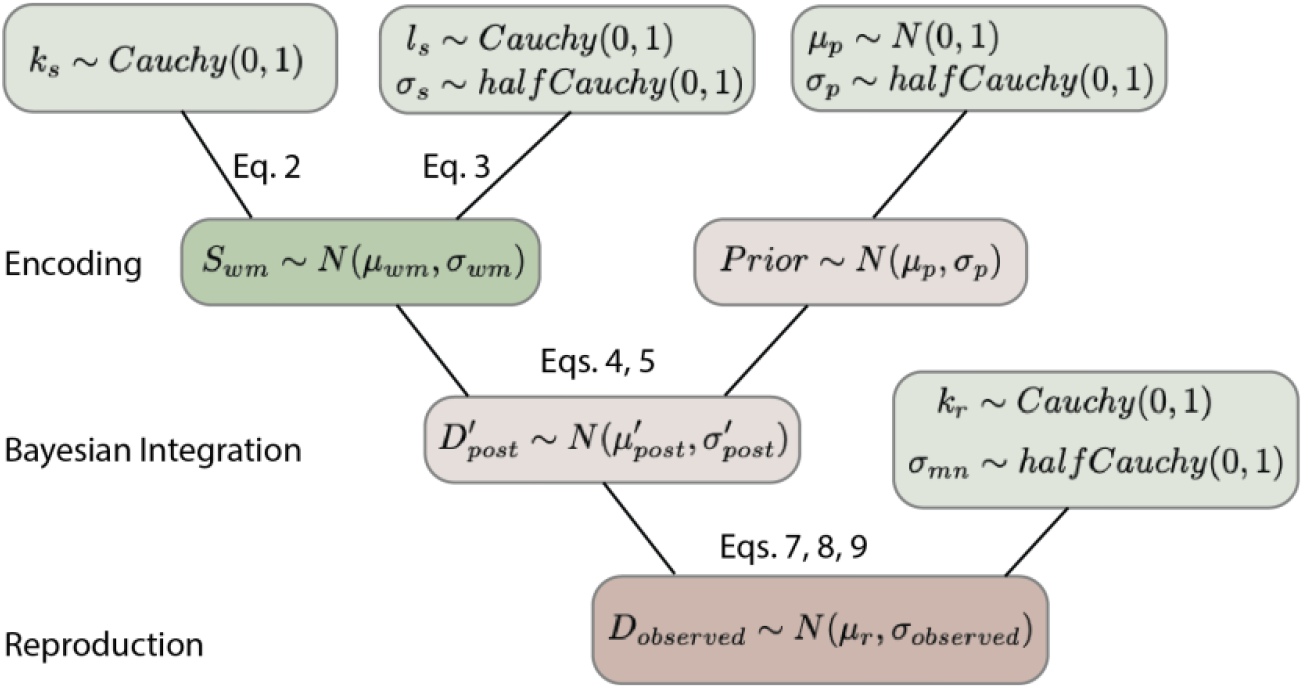
Schematic illustration of the Bayesian hierarchical model through RStan in parameter estimation. Light green boxes show the standard distributions used for the hyperparameters. The green box represents the duration encoding stage, light brown boxes Bayesian integration stage, and the brown box reproduction stage.

After compiling the model specification from Stan’s probabilistic programming language to a C++ program, we used the Markov Chain Monte Carlo (MCMC) sampling method in RStan with 8000 iterations per chain (total of four chains) to estimate the parameters for individual participants by maximizing the joint posterior distribution of parameters of interest. The estimation was performed using the R-package RStan (Carpenter et al., 2017; Stan Development Team, 2018), with the data and R-code available at https://github.com/msenselab/working_memory_reproduction.

## Results

### Memory task

The mean accuracy of individual participants for the working memory test was calculated by the proportion of correct responses, including the ‘hit’ responses (for trials with the same color of the probe and the memory item) and the ‘correct rejection’ responses (for the trial with different colors for the probe and memory item). Figure 3 shows the approximate linear relation between memory load and the mean accuracy in the memory test across the four experiments, with an overall decrease of the correct rates from Experiment 1 to 4.

**Figure 3.**
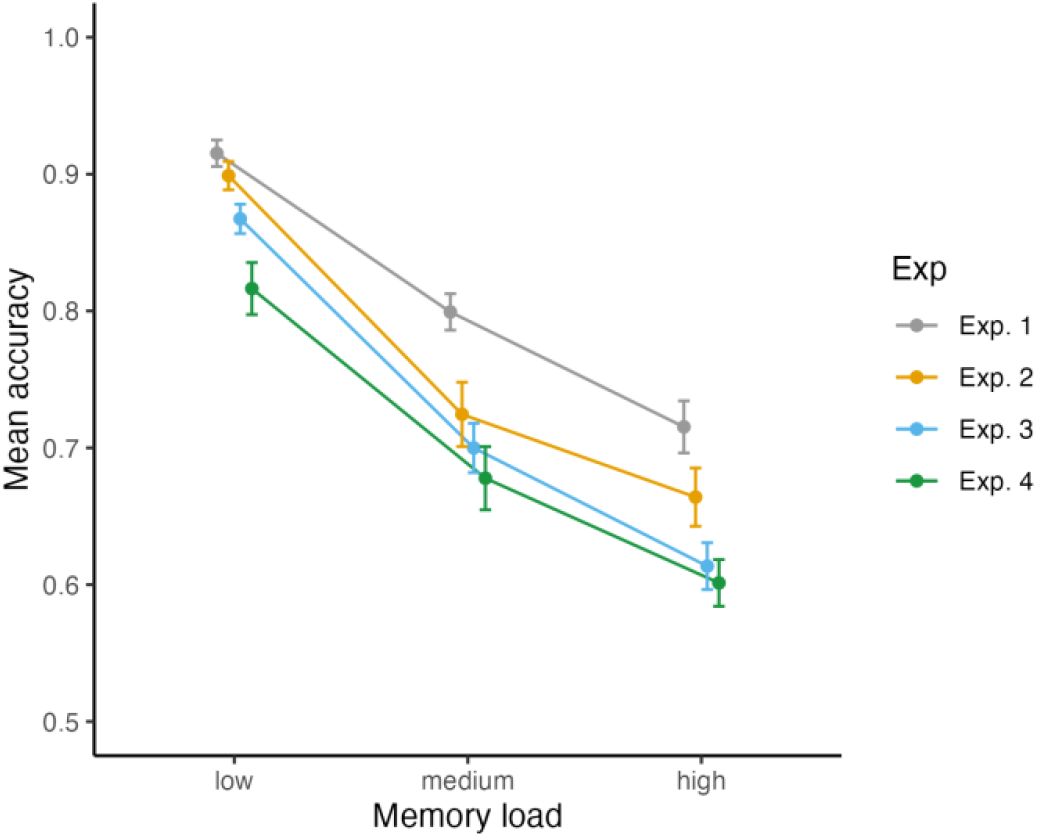
Mean accuracy with associated standard errors (n=16) for the (secondary) workin-gmemory task as a function of the memory load (low, medium, high), separated for Experiments 1-4.

A mixed-design ANOVA of the mean correct rates with Experiment (1–4) as between-subject factor and Memory Load (small, medium and high) as within-subject factor revealed a significant main effects of Experiment, *F*(3, 60) = 11.15, *p* < .001, *η_g_^2^* = .36, and of Memory Load, *F*(2, 120) = 454.22, *p* < .001, *η_g_^2^* = .88. However, the Experiment × Memory-Load interaction did not reach significance, *F*(6, 120) = 2.04, *p* = .066, *η_g_^2^* = .093. The mean accuracy in the working-memory task decreased linearly as the memory load increased from the 1 to 5 items (linear effect, *t(30)*=-*29.181*, *p* < .001, mean of .87, .73, .65 of low, medium and high load respectively), indicating that the manipulation of memory load was effective: larger set sizes engendered higher loads. Further LSD comparisons revealed memory accuracy to be significantly higher in Experiment 1 (mean of .81) than in Experiments 2–4 (means of .76, .73, and .70, respectively; all *p*s < .025), indicating that the working-memory performance declined significantly when the memory task was intermixed with the duration task (in Experiments 2–4). Interestingly, performance was also significantly better in Experiment 2 than in Experiment 4 (*p* < .001), that is: when the memory task was performed first (i.e., when it overlapped with the duration-production phase) vs. second (i.e., when it overlapped with the reproduction as well as the production phase).

### Duration task

Figure 4 shows the reproduction biases and coefficients of variation (CVs) for all four experiments. The CV is a standardized measure of dispersion of reproduced durations (i.e., normalizing the standard deviation by the duration), which is a close approximation to the Weber fraction. As can be seen, there was a central-tendency effect in all experiments, with the short intervals being overestimated and long intervals underestimated, and the CVs decreased as the tested (i.e., to-be-reproduced) durations increased.

**Figure 4.**
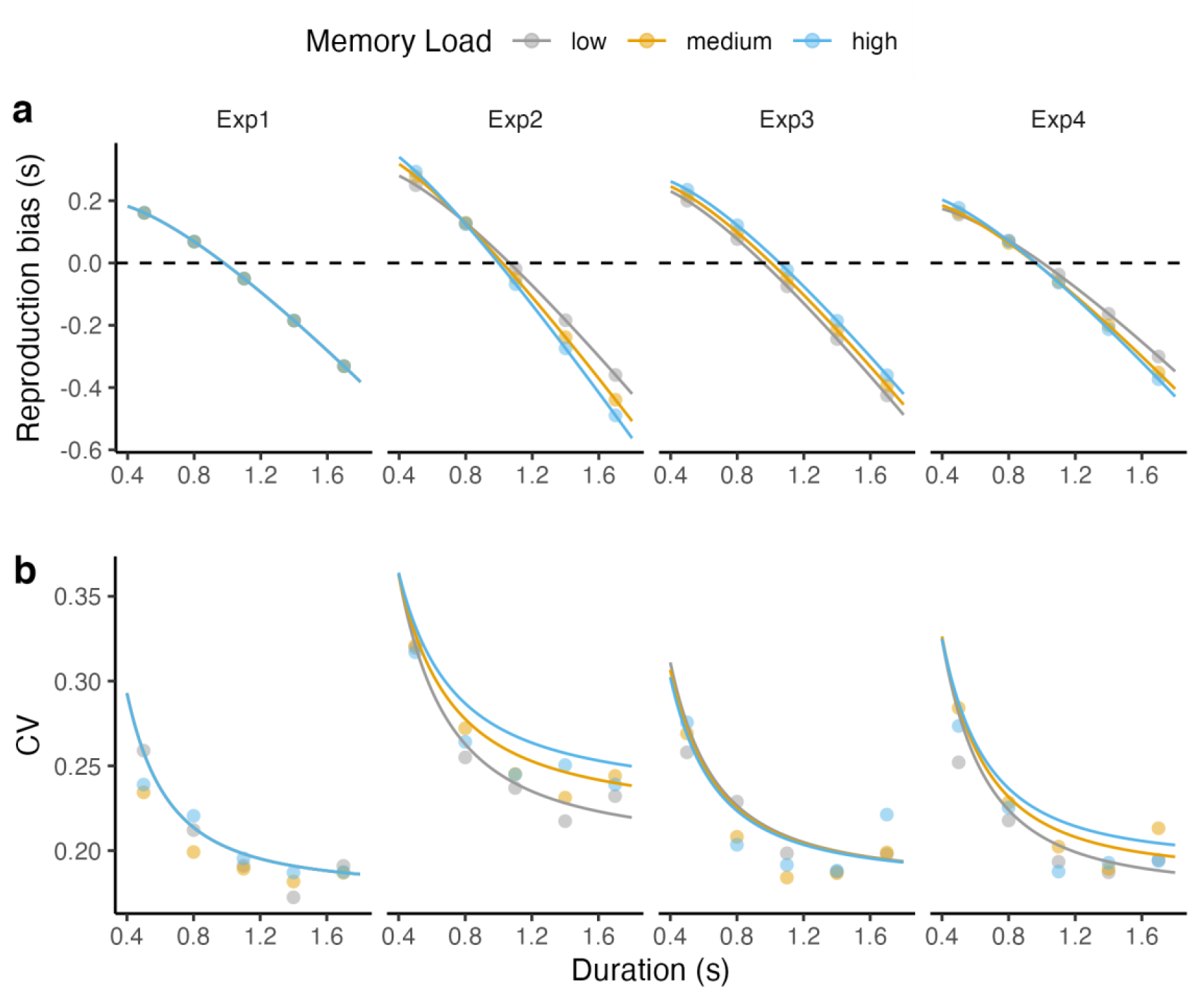
(**a**) Mean reproduction biases and **(b**) coefficients of variation (CVs), for the four experiments. The dots represent the observed mean data, the curves the predictions from the Bayesian model outlined in the modeling section; the gray, orange, and cyan colors represent the low, medium, and high memory-load conditions, respectively.

**Figure 5.**
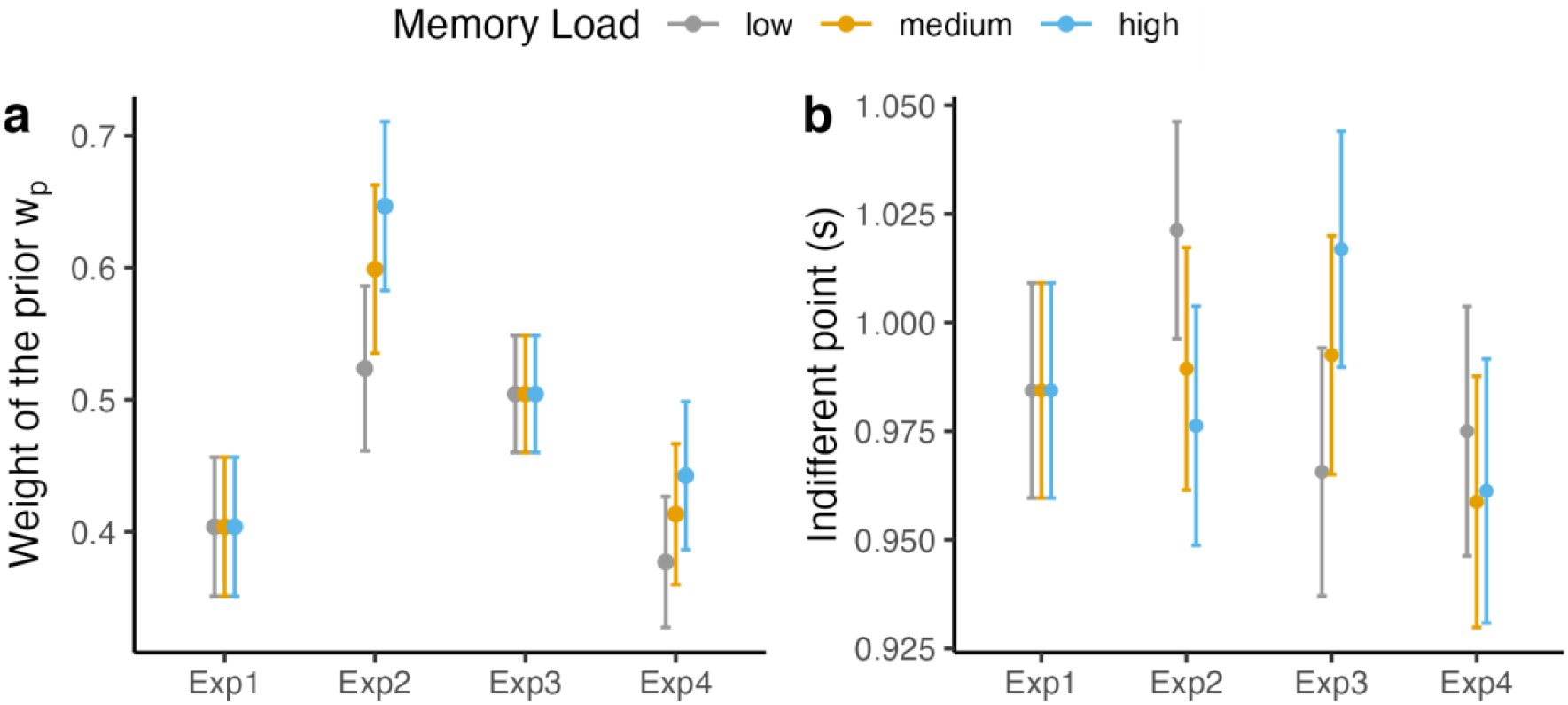
(**a)** Mean central-tendency indices measured by the estimated weight of the prior (w_p_), separately for the three memory-load conditions in the four experiments. (**b)** Mean Indifferent Points, separately for the three memory-load conditions in the four experiments. The gray, orange, and cyan colors represent the low, medium, and high memory-load conditions, respectively. Note, w_p_ was set to the same value for Exps. 1 and 3.

#### The central tendency effects

The central-tendency indices, calculated as the weight of the prior *w_p_* in Bayesian modeling (Figure 4a), were significantly larger than 0 in all four experiments (all w_*p*_s > 0.36, all *t*s > 8.14, all *p*s < 0.001), confirming robust central-tendency effects in our duration estimation tasks.

Because the working-memory task was introduced after the production phase in Experiments 1 and 3, the model assumed no influence of memory load on the central tendency (see Eq. 4 and Figure 3a). The mean weight of the prior (± standard error, SE) was 0.40 ± 0.05 in Experiment 1 and 0.50 ± 0.04 in Experiment 3. Interestingly, the model was able to predict the differential central-tendency effects between Experiments 2 and 4 (see the goodness of fit in Figure 3, and Appendix A). Repeated-measures ANOVAs and tests of the linear relation of the central-tendency indices (*w_p_*) to the factor Memory Load (low, medium, high), conducted separately for Experiments 2 and 4, revealed a significant linear increase of the central tendency with increasing memory load in both Experiment 2 [mean central tendency for the low, median, and high memory loads: 0.52 ± 0.06, 0.6 ± 0.06, 0.65 ± 0.06, respectively; ANOVA: *F*(1.01, 15.2) = 55.09, *p* < .001, *η_p_^2^* =.79; linear effect, *t (30)*= 10.41 *p* < .001] and Experiment 4 [mean central tendency: 0.38 ± 0.05, 0.41 ± 0.05, and 0.44 ± 0.05, respectively; ANOVA: *F*(1.01,15.2) = 33.98, *p* < .001, *η_p_^2^* = .7; linear effect: *t(30)* = 8.23, *p* < .001]. Recall that both Experiments 2 and 4 involved working-memory pressure in the duration-production phase, and this in turn caused stronger central-tendency effects for the high vs. the low memory loads.

When collapsing all levels of memory loads together for individual experiments and comparing across experiments, the mean central-tendency indices turned out to differ significantly among the four experiments [one-way ANOVA: *F*(3, 60) = 2.87, *p* = .044]. Further LSD comparisons revealed significantly higher central-tendency indices in Experiment 2 (.590 ± .061) than in Experiment 1 (0.40 ± .05) and 4 (0.41 ± .04), all *p*s < .05; no other comparisons reached significance. The relatively stronger central-tendency effect in Experiment 2 may be caused by the increased sensory noise in having to perform the memory test task between the production and reproduction phases: this could increase sensory noise either due to the prolonged production-reproduction phase interval or the attentional sharing between the two tasks.

In a further, Wilcoxon-test analysis, we compared the model factor *l_s_* (see Eq. 4), indicative of the extent to which the variance of the sensory measure is modulated by memory load, between Experiments 2 and 4: *l_s_* turned out to be significantly higher in Experiment 2 [mean of 0.201 ± 0.139] vs. Experiment 4 [0.013 ± 0.004; W = 217, p <.001]. That is, sensory noise was larger when the working-memory task overlapped only with the production phase of the duration task (in Experiment 2) as compared to when it overlapped with both the production and reproduction phases (in Experiment 4). In turn, the larger sensory noise led to a higher central-tendency effect in Experiment 2 than in Experiment 4.

#### The mean indifferent points (IP) and general reproduction bias

The mean Indifferent Points (IPs; shown in Figure 4b), at which the duration reproduction is veridical, can be used to index participants’ general reproduction bias: an IP larger than the mean tested duration is indicative of a general overestimate, while a smaller IP is indicative of a general underestimate. A mixed-design ANOVA with Memory Load (low, medium and high) as within-subject factor, Experiment (1–4) as between-subject factor revealed the Load × Experiment to be significant [*F*(6, 120) = 6.110, *p* < .001, *η_p_^2^* = .23], while the main effects were non-significant [Memory Load: *F*(1.01, 40.69) = .429, *p* = .652, *η_p_^2^* = .007; Experiment: *F*(3, 60) = .285, *p* = .836, *η_p_^2^* = .014]. To understand the interaction effect, we examined the Load effect on the IPs separately for each experiment. In Experiment 1, the IPs were essentially the same for the three memory-load conditions (.984 ± .024; see also Figure 3a), given that the working-memory task was introduced after the completion of the duration task. In Experiment 2, the IPs *decreased* linearly with increasing memory load [one-way ANOVA: *F*(1.09, 16.37) = 5.83, *p* = .007, *η_p_^2^* = .28, linear effect: *t(30)* = −3.32, p = .002, means of 1021 ± 26 ms, 989 ± 26 ms, and 976 ± 27 ms for the low, median, and high memory-load conditions, respectively], indicating that a memory load imposed on the production phase caused a significant underestimation of the duration. In Experiment 3, the IPs increased linearly with increasing memory load [*F*(1.03, 15.48) = 8.75, *p* = .007, *η_p_^2^* = .37, linear effect: t(30) = 4.182, p < .001; means of 966 ± 28 ms, 993 ± 27 ms, 1017 ± 26 ms, respectively]. The pattern was opposite to that in Experiment 2, showing that a memory load imposed on the reproduction phase led to an overestimation of the tested duration. In contrast, when the memory load spanned both the duration-production and reproduction phases in Experiment 4, the IPs were comparable among three different memory-load levels [one way ANOVA: *F*(1.04, 15.62) = .74, *p* = .49, *η_p_^2^* = .05; means of 975 ± 28 ms, 959 ± 28 ms and 961 ± 29 ms, respectively]. The comparable IPs among three memory load conditions suggests that the opposite effects of Memory Load observed in Experiments 2 and 3 cancel each other out when the memory tasks spans the whole (production plus reproduction phases of the) duration task.

Recall that participants’ general biases were modeled by two scaling factors, *k_s_* and *k_r_*: *k_s_* represents the magnitude of the duration shortening per unit of memory load in the production phase, and *k_r_* the magnitude of the duration lengthening per unit of memory load in the reproduction phase. When compared to zero, one sample t-tests revealed a significantly positive *k_s_*-value in Experiment 2 (*k_s_* = .263 ± .077, *t* = 3.409, *p* = .004), a significantly positive *k_r_*-value in Experiment 3 (*k_r_* = .025 ± .003, *t* = 7.212, *p* < .001), and both a significantly positive Æ_s_-value and a significantly positive *k_r_*-value in Experiment 4 (*k_s_* = .459 ± .052, *t* = 8.824, *p* < .001; *k_r_* = .229 ± .047, *t* = 4.924, *p* < .001). These results are confirmatory of the attentional-sharing hypothesis, that is: the concurrent memory and duration estimation tasks share the same attentional resource, which in turn leads to a duration underestimation with working memory pressure during the production phase, but duration overestimation with memory pressure during the reproduction phase. When the memory pressure occurs in both phases, the duration underestimation in production and overestimation in reproduction may cancel out each other, resulting in the diminishing of the general bias.

#### Coefficient of variations of the duration reproduction

The observed Coefficients of Variation (CVs) for each reproduced sample duration, calculated as *CV_i_* = *σ_i_*/*R_i_*, where *σ_i_* and *R_i_* represent the standard deviation and the mean of the reproduction of a given interval *D_i_*, are shown in Figure 4b. A mixed-design ANOVA of the CVs, with Sampled Duration (500, 800, 1100, 1400, 1700 ms) and Memory Load (low, medium, high) as within-subject factor and Experiment as between-subject factor revealed only the main effects of Duration and Experiment to be significant; no other effects (including the main effect of memory load and all interactions) reached significance (all Fs < 1.47, all ps > .19). The main effect of Duration (*F*(2.429, 7.286) = 47.794, *p* < .001, *η_p_^2^* = .44) was due to the CVs decreasing linearly with increasing sample duration (means of .274 ± .008, .229 ± . 006, .205 ± .005, .195 ± .005, and .209 ± .006 from short to long durations, respectively; linear effect: t(60) = −11.423, p < .001]. And the main effect of Experiment (*F*(3, 60) = 6.535, *p* = .001, *η_p_^2^* = .246; means of .203 ± .010, .259 ± .010, .213 ± .010, and 216 ± .010 in Experiment 1–4, respectively] was owing to the significantly higher CV in Experiment 2 compared to the other experiments (LSD tests comparing Experiment 2 with Experiment 1, 3, and 4, respectively: all ps < .003; comparable CVs in Experiment 1, 3, and 4). The relatively larger response variation in Experiment 2 is likely due to the working-memory test phase being administered in-between the duration-production and reproduction phases, while in all the other three experiments the memory task was tested at the end of the reproduction phase.

#### Goodness of the modeling

The Bayesian model outlined above predicted the behavioral results strikingly well, in all four experiments. The model performance may be gauged by calculating the means of the relative prediction error on the estimated durations(*Mean*(|*D_observed_* – *D_predicted_*|/*D_observed_*)) and on the estimated variances (*Mean*(|*SD*(*D_observed_*) – *sd*(*D_predicted_*)|/*sd*(*D_observed_*))). The results revealed less than 4.17% error on the reproduced durations for all experiments (3.34%, 4.17%, 3.44%, and 3.53% for Exp. 1–4 respectively; see Appendix A for further details about the goodness of the model fitting).

## General Discussion

The present study examined cognitive-load interference in duration estimation using a Bayesian approach. Through computational modeling of the results from four experiments, we attempted to provide a generative model of load interference on the whole champion of processes in duration estimation, including duration encoding and reproduction. The Bayesian model we proposed predicted not only the mean but also the coefficient of variation (CV) of reproduction behavior.

We found a visual working-memory load to interfere with participants’ duration reproduction both when the load was imposed during the duration-production and during the reproduction phase, albeit in a different way. In more detail, when the working-memory task overlapped only with the production phase (in Exp. 2), participants on average underestimated the tested durations, and they exhibited a stronger central-tendency effect under high- vs. low memory-load conditions. In contrast, when the working-memory task overlapped only with the reproduction phase (in Exp. 3), the higher the memory loaded, the more duration participants over-reproduced, while the central-tendency effect was comparable across the different load conditions. Of note, when the workin-gmemory task spanned both the production and reproduction phases (in Exp. 4), there was no longer a general over- or underestimation, but the central-tendency effect remained stronger with higher vs. lower memory loads. Finally, varying levels of memory load introduced between consecutive duration reproductions (in Exp. 1) had no discernible effects on either general reproduction biases (i.e., there was no general over-/underestimation) or the central-tendency bias, suggesting that the prior updating of the sampled durations was not influenced by the intermediate secondary task.

Importantly, this pattern of findings could be well fitted by our Bayesian model, which integrated the notion of attentional-resource (Fortin, 2003; Fortin & Rousseau, 1998; Macar et al., 1994) between two concurrent tasks. According to the Bayesian inference model (Jazayeri & Shadlen, 2010; Petzschner et al., 2015; Shi et al., 2013), the reproduced duration reflects an optimal integration (according to the Bayes rule) of the sensory estimate of a given duration with the prior distribution stored in the memory, where ‘optimal’ refers to achieving minimal variability in the final estimate. A by-product of this is that the duration estimates assimilated to the mean prior, as evidenced in the typical central-tendency bias. Standard Bayesian inference models make no assumptions about how memory load may influence Bayesian inference. To address this, here we combined the attentional-sharing account (Fortin, 2003; Fortin & Rousseau, 1998; Macar et al., 1994) with standard Bayesian inference, assuming that time units would be lost in the duration encoding and reproduction stages when attention is shared with a secondary task. Prior work (Fortin, 2003; Fortin & Rousseau, 1998) had shown that the loss of time units in the encoding and reproduction stages has a differential impacts: when attention is diverted away from the primary (temporal) task by another, concurrent non-temporal task during the duration-encoding phase, a certain amount of clock ticks would be lost, resulting in a shortened time estimation (underestimation); in contrast, when the secondary task is performed concurrently with the reproduction phase, the resulting loss of clock ticks (due to lapses in monitoring the elapsed time) would lead to a reproduced duration longer than the tested interval (overestimation). This descriptive explanation is quantitatively characterized by the linear scaling parameters *k_s_* and *k_r_* in our Bayesian model (Eqs. 2 and 7). Both the behavioral findings and the model confirm the dissociable influences of memory pressure at different stages of time estimation. Specifically, the concurrent working-memory task gave rise to an underestimation when imposed during the production phase (*k_s_* > 0), but an overestimation when imposed during the reproduction phase (*k_r_* > 0).

While the attentional-sharing account (Fortin, 2003; Fortin & Rousseau, 1998; Macar et al., 1994) can well explain the general over- and, respectively, underestimation of the sample durations, it falls short in explaining the differential central-tendency effects. While there were central-tendency effects in all four experiments, the central-tendency bias was significantly modulated by memory load only in Experiments 2 and 4, that is, when the secondary memory task overlapped the production phase. We take this to indicate that the central-tendency bias was introduced mainly in the duration-encoding stage: the memory load imposed during this stage increased the uncertainty of the sensory measure (which we modeled with the scaling parameter *l_s_* in Eq. 3), translating into a reduction of the sensory weight in Bayesian integration and, in turn, an increased central-tendency bias. In contrast, making the secondary task overlap the reproduction phase (Exp. 3) or introducing it after completion of the reproduction (Exp. 1) did not significantly change the central-tendency effect, which corroborates that the Bayesian integration occurred primarily at the encoding stage.

It should be noted that the influence of the memory load on the central tendency was more pronounced in Experiment 2 than in Experiment 4, where, in the latter, the secondary task extended across both the duration-encoding and reproduction phases of the primary task. In the model, this differential load effect was captured by the value of *l_s_* – the scaling parameter of the sensory-measurement uncertainty – being significantly larger in Experiment 2 vs. Experiment 4. This finding raises a question: if the modulation of the central-tendency effect exclusively arises in the encoding stage, shouldn’t the two experiments have engendered comparable effects of the memory load on the central-tendency bias? The fact that they didn’t might be explained by the order of the primary and secondary tasks. In Experiment 2, the secondary memory task was the first task requiring a response (i.e., the memory test preceded the reproduction), whereas it was the second task in Experiment 4. As can be seen from Figure 2, accuracy in the memory test was significantly higher in Experiment 2 than in Experiment 4, that is, when the memory was probed first rather than second. Within the attentional-sharing framework, this pattern would indicate that more attentional resources were allocated to the first than to the second task. Consequently, allocating relatively less attention to the duration task in Experiment 2 relative to Experiment 4 would have led to an increase of the uncertainty of the duration estimates, rendering a stronger central-tendency effect. In addition, probing the secondary memory task first also lengthened the gap between the encoding and reproduction stages, which might cause additional forgetting of the estimated duration. Such forgetting might assimilate the representation toward the distal (i.e., long-term) mean prior, as would be suggested by the adaptation-level theory (Helson, 1964). The influence of the prolonged gap between the encoding and reproduction stages was also numerically, though not significantly, evident in Experiment 3 as compared to the baseline Experiment 1 (see Fig. 4a). Unfortunately, because we did not record the completion time of the memory test in between the production and reproduction phases, we cannot quantitatively determine the impact of the prolonged gap on the central tendency bias. Thus, this conjecture deserves further investigation in future studies.

Interestingly, (Allred et al., 2016) recently reported in a line-length judgment task that the central tendency is likewise influenced by the memory load. In their study, the memory items had to be held in working memory for the whole process of the primary line-judgment task, which is similar to our Experiment 4. The consistent influences of memory load on the central-tendency effect in non-temporal (Allred et al., 2016) as well as temporal tasks (the present study) suggest that the Bayesian model we propose here is generic, rather than being limited to the time domain. Given that Allred et al.’s study design did not separately manipulate the memory load in the encoding and reproduction stages, their finding does not tell at which stage the interference occurred. Here, with four experiments imposing the memory load in different stages, we found that the impact on the central-tendency effect was primarily attributable to cognitive-load interference during the encoding, rather than the reproduction (retrieval), stage – an attribution that informed the construction of our computational model.

As briefly discussed earlier, when the secondary memory task was imposed on the reproduction phase in Experiment 3, the central tendency was not influenced by the memory load. Interestingly, though, we saw a general shift (bias) in the reproduced duration (Figs. 3a): the higher the memory load, the larger a (general) shift we observed in the reproduced durations – as could also be seen in the shift of the indifference points (Fig. 4b). The dissociation between the central tendency and the general bias mainly came from the differential interference of the memory load in the duration-encoding and reproduction stages. The reproduction stage did not involve any Bayesian integration, just comparing the elapsed time to the estimated duration retained in memory. The primary impact of the memory load consists of the lapse of attention in monitoring the elapsed time (Fortin & Rousseau, 1998; Glasauer et al., 2007), which causes loss of some units of passage time and thus an over-reproduction of the (estimated) duration. This would explain the general shift in the reproduced duration that we observed (captured by Eq. 7, also Eq. 10). Interestingly, the varying memory loads did not alter the memory representation of the reference (i.e., the estimated) duration, which was derived by Bayesian integration in the encoding stage. However, as considered above, across experiments the reference duration might have been influenced by the temporal gap between the encoding and reproduction stages. The fact that the memory load failed to change the reference duration suggests that the representation of the single reference duration is rather robust. However, this might change if the task requires two or more reference durations to be held in memory – a conjecture that would be interesting to examine in a future study.

It should be noted that the general shift in the reproduced duration was not limited to Experiment 3: we also observed a general shift in Experiment 2, though in the opposite direction (Fig. 4b). As shown in model Eq. 10, unlike the central-tendency effect (captured by *w_p_*) which is only influenced by the memory load in the encoding stage, the indifference point is influenced by both stages, though in opposite directions. When a load was imposed only on the encoding stage, underestimated durations with higher vs. lower loads caused a general downshift in the indifference points, in addition to the influence on the central-tendency bias in the encoding stage. Given the opposite influences of memory load in the encoding and reproduction stages, we observed the opposite trends in Experiments 2 and 3. This could then also explain the absence of significant shifts in Experiment 4, as the net impacts of the general shifts roughly canceled each other out when the memory load was imposed on both stages (in Experiment 4).

In summary, imposing the memory load on the encoding and reproduction stages of duration estimation, we replicated the general underestimation and overestimation of a given duration when the memory load was increased in the encoding and reproduction stages, respectively, as suggested by the attentional-sharing account. In addition, we found the central-tendency bias was only influenced by the memory pressure in the encoding stage. Using a generative Bayesian model, we detailed when and how memory pressure affects time estimation and the concomitant effects on the central-tendency bias, and quantitatively predicted behavioral results from all four experiments. Last but not the least, the generative model we proposed here for the influence of the cognitive load on time perception might be generalizable to other forms of magnitude perception.

## Appendix

## Appendix A. Model comparison

The three-stage Bayesian model introduced here assumes logarithmic encoding of subjective durations, based on Fenchner’s logarithmic law (Fechner, 1860). Logarithmic encoding implicitly assumes that time percepts follow the scalar property (Gibbon et al., 1984; Shi et al., 2013), namely, a constant of Weber fraction of time estimation. It should be noted that it is not the only model in the field. An alternative assumption holds that the internal representation is linearly scaled, but with the noise increasing linearly with the absolute magnitude according to Weber’s law. However, empirical justification of linear vs. logarithmic coding of the internal representation largely depends on the adopted experimental paradigms (Maaß et al., 2021; Matthews & Meck, 2016; Ren et al., 2021). Accordingly, we explored both logarithmic and linear encoding models and compared their predictions to determine which model performs better.

In the logarithmic-encoding model, the duration is first transformed to the logarithmic scale. Bayesian integration of the sensory input and memory prior, and the influences of the memory load operate on this scale. The duration thus estimated is then transformed back to the linear scale for the reproduction. The memory influence occurs at the reproduction stage on the linear scale (see main text for details of the model). In contrast, the linear-encoding model assumes that all processes operate at the linear scale. However, the model further assumes Weber scaling, that is: the sensory measure (S) of given sample duration (D) follows Weber’ law: *S* = *D* + *ϵ*, where *ϵ* indicates internal measurement noise. The standard deviation of sensory measurement (*σ_s_*, estimated from the noise *ϵ*) is approximately proportional to the mean of the sensory measurement (*μ_s_*), *σ_s_* =*k*\* *μ_s_*, where *k* is known as the Weber fraction of sensory measurements. Similar to the logarithmic-encoding model, both the mean estimate and its standard deviation are assumed to be linearly affected by the memory load. Let the memory representation without loss of clock ticks be normally distributed, 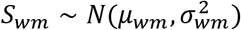. Accordingly, both the mean *μ_wm_* and the variance 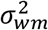 are subject to the influence of the memory load:

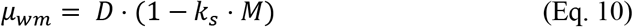

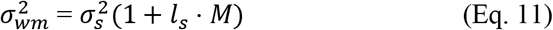

At the Bayesian integration stage, the distribution of the internal prior is assumed as 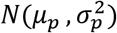 with the mean *μ_p_* and the variance 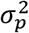, and the prior is integrated with the memory representation of duration *S_wm_* according to the Bayesian rule, so that Eq. 4 and Eq. 5 are also applicable in the linear-scale model. During the reproduction process, time units could be lost due to attentional sharing and reproduction inherited additional motor noise. Let 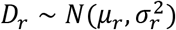 be the reproduction distribution with:

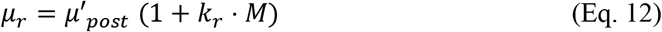

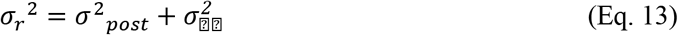

The notations of these key parameters in the linear model are the same as those in the logarithmic model.

Both the logarithmic- and the linear-scale model perform well in predicting the mean reproduction and the coefficient of variation (CV). However, while the predictions are comparable as regards the mean reproduction, the logarithmic model predicted the CV significantly better than the linear model. Figure A1 represents the mean absolute errors (MAE) of the predictions derived from the linear- and, respectively, logarithmic-scale models (reproduction and CV): each point represents the mean absolute prediction error in the reproduction means and their CVs in the various conditions across the four experiments. As can be seen, the logarithmic model produced generally smaller prediction errors than the linear model, with a particularly marked advantage in the CVs. The prediction errors of the logarithmic model never exceed 0.0155, indicated by the dashed line; but the linear model performed worse for more than half the conditions compared to the poorest condition from the logarithmic model.

To formally evaluate the models’ performance, we calculated Watanabe–Akaike information criterion (WAIC) and leave-one-out cross-validation (LOO-CV) as predictive information criteria for Bayesian models, using the Loo package in R framework (Vehtari et al., 2017; Watanabe & Opper, 2010). Lower values of WAIC and LOO-CV imply higher prediction accuracy. Table A1 lists the averaged WAICs, LOO-CVs, and prediction accuracies (AUC) of the reproduction means and variances across all participants. As can be seen, the logarithmic-scale model was associated with lower WAIC and LOO-CV values and higher prediction accuracies across all experiments than the linear model.

**Figure A1.**
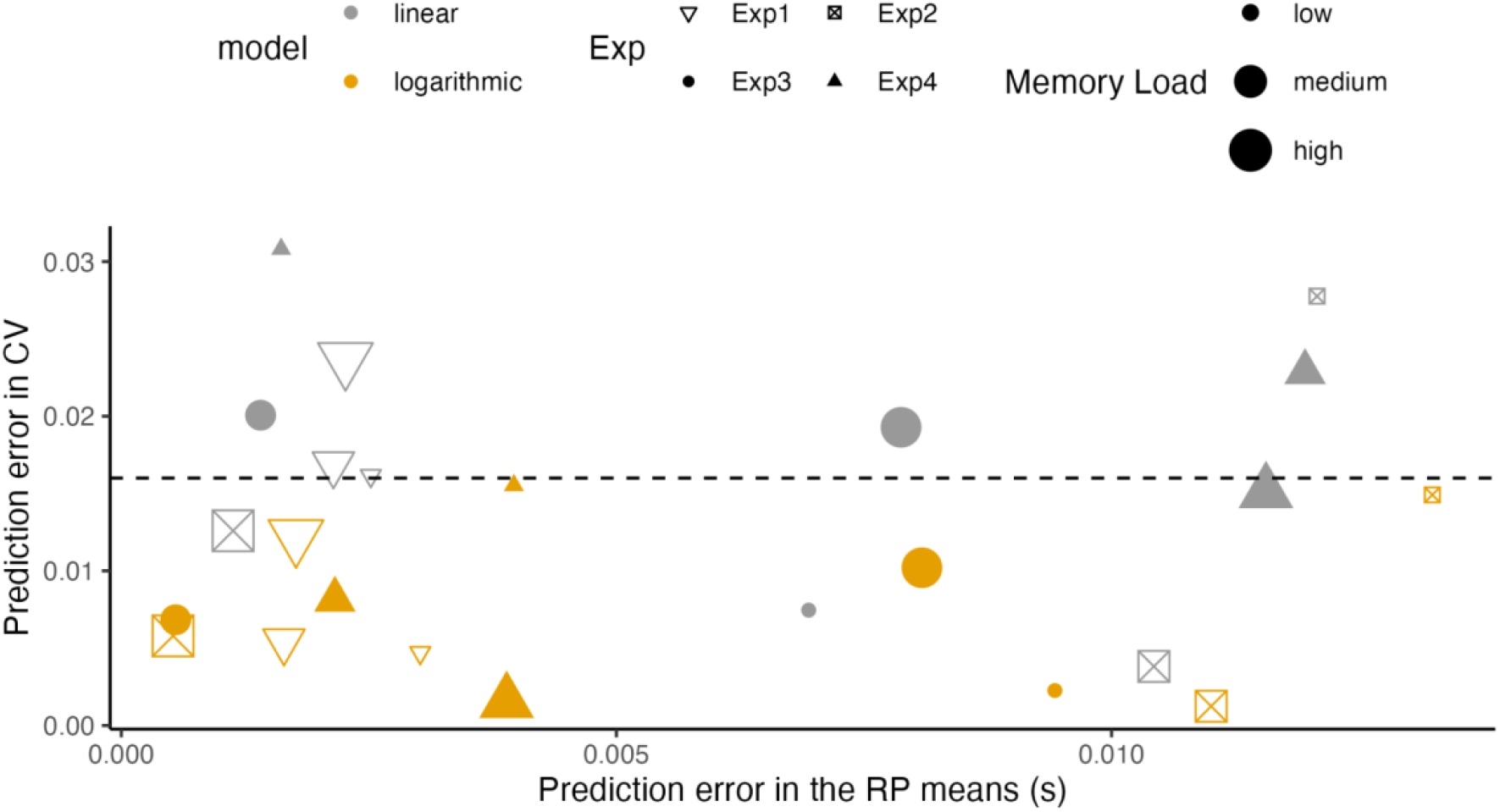
Averaged absolute prediction errors of the reproduction and CV derived from the proposed logarithmic- and linear-scale models for individual observers in the four experiments.

**Table A1.**
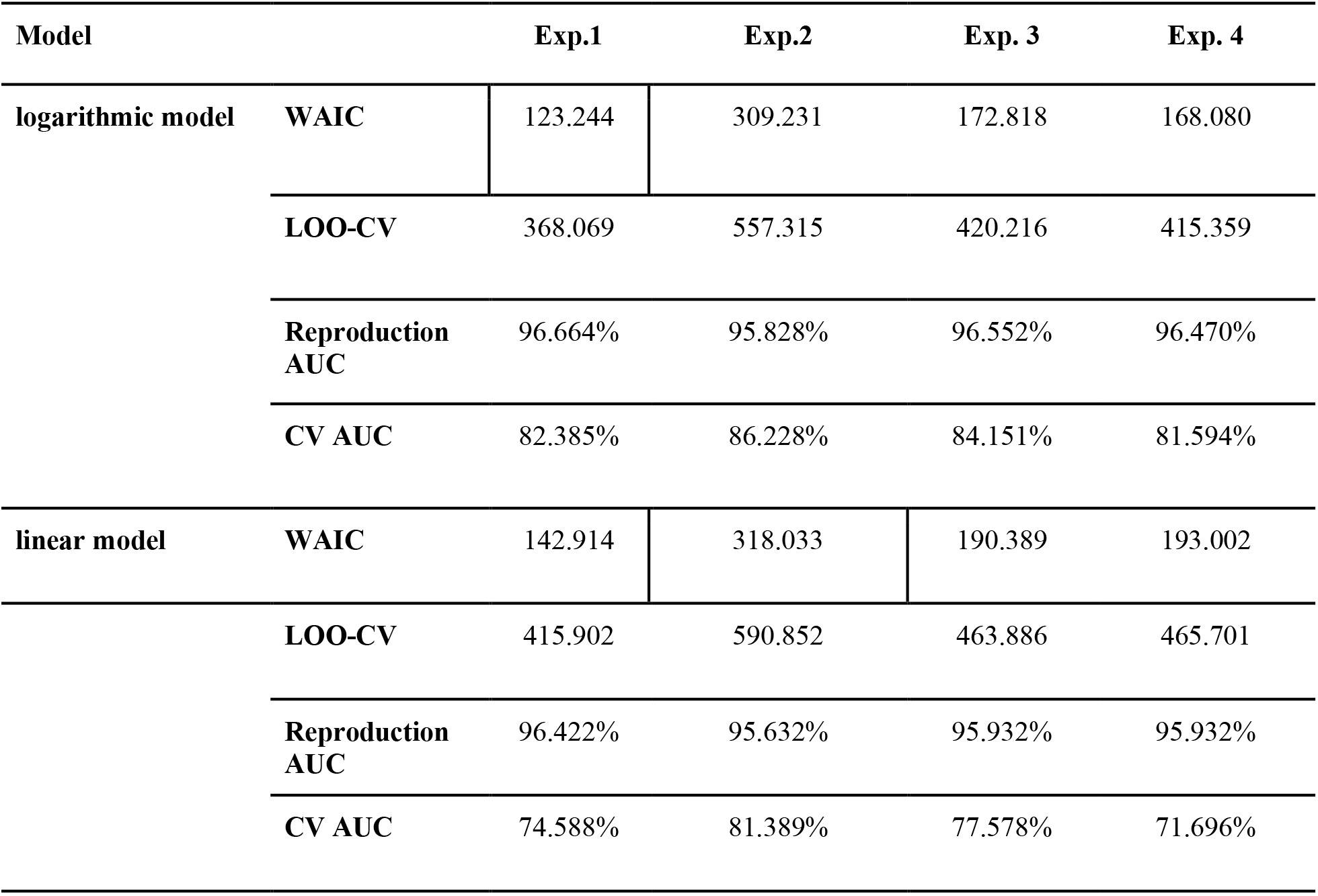
WAIC and WBIC as predictive information criteria for Bayesian models.

## Footnotes

1 This ‘linearity’ assumption does not rule out that the true relation is more complex. In principle, the linear constraint can be relaxed by assuming independent impacts of the memory load on the memory representation. However, such an approach has more free parameters than the current model, and it does not provide additional insights of the underlying mechanism.

2 In the memory-load task, the set sizes were 1, 3, and 5 items of the low-, medium-, and high-load conditions, respectively. It has been shown that the variance is an approximate linear function of set size (for a review, see Bays, 2015). In addition, performance accuracy in the memory task decreased approximately linearly with the set size (see Figure 2). Accordingly, here we set M as a simple linear function of the set size (M = 2 × Set Size - 1) and assume linear relations in Eqs. 2 and 3, without loss of generality.

3 Suppose the duration reproduction engenders an approximate linear central tendency effect. With simple mathematical calculation, the mean reproduction bias is equivalent to *w_p_* times the difference between the indifference point (IP) and the mean sample duration.

